# Divergent effect of central incretin receptors inhibition in a rat model of sporadic Alzheimer’s disease

**DOI:** 10.1101/2021.08.23.457308

**Authors:** Jelena Osmanovic Barilar, Ana Knezovic, Jan Homolak, Ana Babic Perhoc, Melita Salkovic-Petrisic

**Affiliations:** Department of Pharmacology and Croatian Institute for Brain Research, School of Medicine University of Zagreb, 10000 Zagreb, Croatia

**Keywords:** glucagon-like peptide-1, gastric inhibitory polypeptide, hippocampus, hypothalamus, streptozotocin

## Abstract

The incretin system is an emerging new field that might provide valuable contributions to the research of both pathophysiology and therapeutic strategies in the treatment of diabetes, obesity, and neurodegenerative disorders. This study aimed to explore the role of central glucagon-like peptide-1 (GLP-1) and gastric inhibitory polypeptide (GIP) on cell metabolism and energy in the brain as well as on the levels of these incretins, insulin and glucose, by inhibiting the central incretins’ receptors following intracerebroventricular administration of the respective antagonists in healthy rats and a streptozotocin-induced rat model of sporadic Alzheimer’s disease (sAD). Chemical ablation of the central GIP receptor (GIPR) or GLP-1 receptor (GLP-1R) in healthy and diseased animals indicated a region-dependent role of incretins in the brain cell energy and metabolism and central incretin-dependent modulation of peripheral hormone secretion, markedly after GIPR inhibition, as well as a dysregulation of the GLP-1 system in experimental sAD.

## Introduction

Alzheimer’s disease (AD) is a progressive neurodegenerative disease closely connected to the brain insulin resistant state and dysregulated glucose metabolism (Kellar and Craft, 2020). A growing body of research shows that diabetes mellitus type 2 (T2DM) is a risk factor for AD. In line with that, drugs that can treat T2DM effectively may also elicit neuroprotective effects (Kellar and Craft, 2020) and the newest trend in AD research are drugs acting on the incretin system (Femminella et al., 2019; Gejl et al., 2016; Hölscher, 2014).

Gastric inhibitory polypeptide (GIP) and glucagon-like peptide-1 (GLP-1) are the most important incretins for glucose regulation secreted by the gut upon food intake. As far as we know, their primary goal is to stimulate insulin secretion from pancreatic β cells (Sandoval and D’Alessio, 2015; Yabe and Seino, 2011). Both incretins are synthesized in the gut starting with secretion of GLP-1 from sweet taste receptors in the tongue accompanied by secretion of GIP from K-cells in the duodenum and jejunum, ending with secretion of GLP-1 from L-cells in the ileum and colon (Mortensen et al., 2003; Seino et al., 2010; Takai et al., 2015). Upon their secretion, the incretins are degraded by dipeptidyl peptidase 4 (DPP-4) which has a higher affinity towards GLP-1 than GIP. The most accepted physiological role of GLP-1 and GIP upon secretion, is stimulation of insulin secretion by binding to the GIP receptor (GIPR) or the GLP-1 receptor (GLP-1R) on the pancreas β cell (Seino et al., 2010). These receptors belong to the G-protein coupled receptor family, and activate adenylate cyclase and increase intracellular cAMP levels in pancreatic β cells, thereby stimulating insulin secretion in a glucose-dependent fashion (Baggio and Drucker, 2007; Yabe and Seino, 2011). Additionally, GLP-1 slows down gastric emptying and inhibits glucose-dependent glucagon secretion. In contrast, GIP promotes energy storage via direct actions on adipose tissue and stimulates bone formation via stimulation of osteoblast proliferation and inhibition of apoptosis (Baggio and Drucker, 2007).

Besides target sites on the periphery, these closely connected incretins also have targets in the brain. GLP-1 is produced in the nucleus of the solitary tract and the caudal brainstem, which project to the hypothalamus and cortical, hypothalamic, and hippocampal nuclei (Rinaman, 2010), while opposite findings were discovered regarding GIP production in the brain (Adriaenssens et al., 2020; Kaneko et al., 2019; Tseng et al., 1993). GIP and GLP-1 receptors are expressed in the cerebral cortex, hippocampus, olfactory bulb, mammillary bodies, brain stem, area postrema and cerebellum (Adriaenssens et al., 2020; Gu et al., 2013). GLP-1 promotes satiety and GLP-1 receptor activation is associated with weight loss in both preclinical and clinical studies (Gallwitz, 2009). However, there is more to these hormones than their action on food consumption and weight gain.

Recently, new studies showed that incretins influence several pathways in the brain, including neuroinflammation, mitochondrial function, cellular proliferation and apoptosis (Athauda and Foltynie, 2016). Deficits in cell energy function, neuronal dysfunction and neuroinflammation are known hallmarks of neurodegenerative disorders such as AD (Yin et al., 2016). These features of AD can also be found in certain animal models, like the rat model induced by intracerebroventricular streptozotocin (STZ-icv) (Correia et al., 2011; Salkovic-Petrisic and Hoyer, 2007). Lately, incretin analogues have also been tested as possible therapeutic agents in the STZ-icv rat model of sAD (Li et al., 2020; Paladugu et al., 2021), since STZ is considered a diabetogenic compound. Analogues of incretins showed a neuroprotective effect, reduction in Aβ deposition, decreased inflammatory response, enhancement of synaptic plasticity, hippocampal neurogenesis, long-term potentiation (LTP), prevention of hippocampal synapse loss and oxidative stress and improvement of cognitive deficit in animal AD models (Adriaenssens et al., 2020; Athauda and Foltynie, 2016; Hölscher, 2018; Li et al., 2020; Paladugu et al., 2021).

The underlying mechanism responsible for the beneficial effect of GLP-1 and GIP may include stabilization of the outer mitochondrial membrane, preventing efflux of cytochrome c (CytC) into the cytoplasm and reducing the activation of caspase 9 and 3, subsequently resulting in reduced apoptosis and oxidative stress through activation of cAMP and other kinases downstream of their signaling cascade (Athauda and Foltynie, 2016; Baggio and Drucker, 2007). A growing body of evidence also indicates that AMP-activated protein kinase (AMPK), being an energy sensor, can be activated by GLP-1R agonists. It seems that GLP-1 signaling activates AMPK by phosphorylation of Thr172, leading to increased glucose uptake by the cells in an insulin-independent manner (Andreozzi et al., 2016), implicating another role of incretins in brain energy homeostasis. Newer data also indicate that AMPK can increase the synthesis of GLP-1 in pancreatic β cells (Jiang et al., 2016). Both GIP and GLP-1 can influence pyruvate dehydrogenase (PDH) activity, by increasing aerobic glycolysis, which in turn increases production of pyruvate, the major substrate for PDH. PDH converts pyruvate to acetyl-CoA, which enters the citric acid cycle to produce adenosine triphosphate (ATP) (Milne, 2013). In that way, incretins are shifting the cell metabolism from oxidative phosphorylation and increased ROS production to aerobic glycolysis leading to a neuroprotective effect (Zheng et al., 2021). In contrast to aforementioned animal studies, studies in human sAD are focused on exploring the effect of GLP-1 analogues and dual GIP/GLP-1 analogues on cognition, without exploring the pathophysiological background of their beneficial effect in neurodegeneration (Femminella et al., 2019; Gejl et al., 2016; Hölscher, 2018). As the incretin system is now a compelling drug target, and the knowledge of its role in physiological and pathological processes in the brain are obscure, the aim of this study was to explore more closely the role of central GLP-1 and GIP on cell metabolism and energy in the brain as well as on the levels of these incretins, insulin and glucose, by inhibiting the central incretins’ receptors following intracerebroventricular administration of the respective receptor antagonist. This chemical ablation of central GIPR or GLP-1R in healthy animals and a streptozotocin-induced rat model of sAD has pointed to the region- and incretin-dependent role in the brain cell energy and metabolism, and to the incretin-dependent modulation of the brain-gut axis, as well as to dysregulation of these functions in experimental sAD.

## Results

### Peripheral changes were more pronounced following central GIPR inhibition

Central inhibition of GIPR had a stronger impact than GLP-1R inhibition on plasma levels of the measured hormones (Fig 2A and B). Both healthy and STZ-icv-treated rats demonstrated a significant increase of plasma hormone levels after GIPR inhibition; insulin (118%, p=0.0016 and 73%, p=0.0524), and total (310.9%, p=0.0343 and 137%, p=0.0002) and active GIP (212-fold, p=0.0004) and 96-fold, p<0.0001), respectively (Fig 2B). In contrast, central GLP-1R inhibition induced no acute changes in plasma hormones in healthy rats but decreased active GLP-1 level in the STZ group (−61.3%; p=0.0019) (Fig 2A). Plasma glucose concentration remained unchanged regardless of the treatment in both groups (Plasma glucose concentration in Fig 2A already published in Supplement 2 (Homolak et al., 2021)). Although the effect could have been masked by increased baseline glucose concentration due to icv treatment.

**Fig 1.**
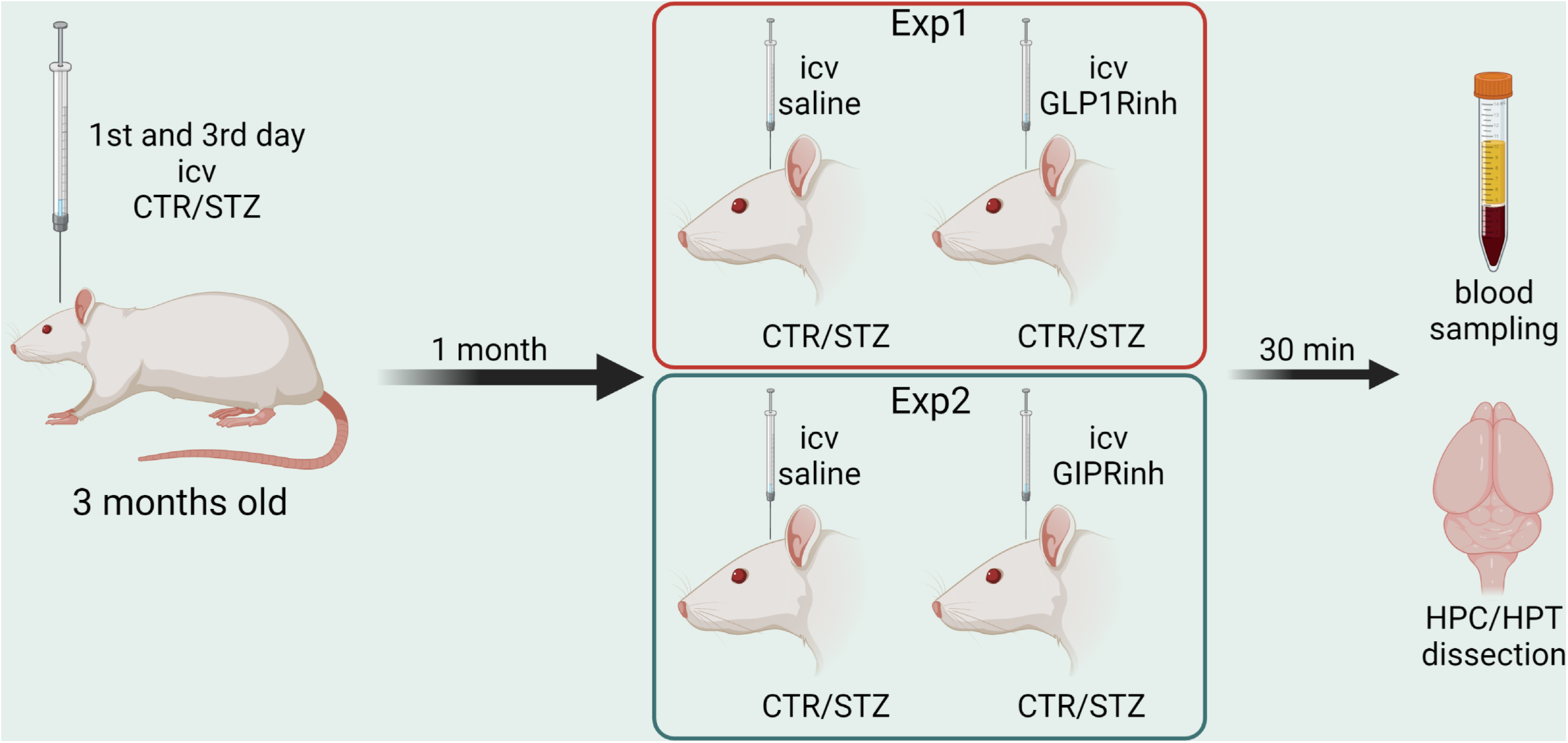
Experimental design. Two experiments with 4 groups each were conducted with the same procedure. Three-month-old male Wistar rats were intracerebroventricularly (icv) injected with streptozotocin (STZ/3 mg/kg) or vehicle only (controls/CTR) on days 1 and 3. In experiment 1 (Exp1) rats were randomly divided into four groups. Two groups (CTR+GLP1Rinh and STZ+GLP1Rinh) received icv 85 µg/kg of glucagon-like peptide-1 receptor antagonist (Exendin fragment 9-39, GLP1Rinh) dissolved in saline. In experiment 2 (Exp2) two groups (CTR+GIPRinh and STZ+GIPRinh) received icv 85 µg/kg of gastric inhibitory polypeptide receptor antagonist (Pro3-GIP, GIPRinh) dissolved in saline. The other groups in both experiments (CTR and STZ) received saline only (bilaterally 2 µL/ventricle). All animals were sacrificed 30 minutes after icv. Blood was sampled and brain was removed and the hippocampus (HPC) and hypothalamus (HPT) were dissected out for further analysis. Figure was created with BioRender.com.

**Fig. 2.**
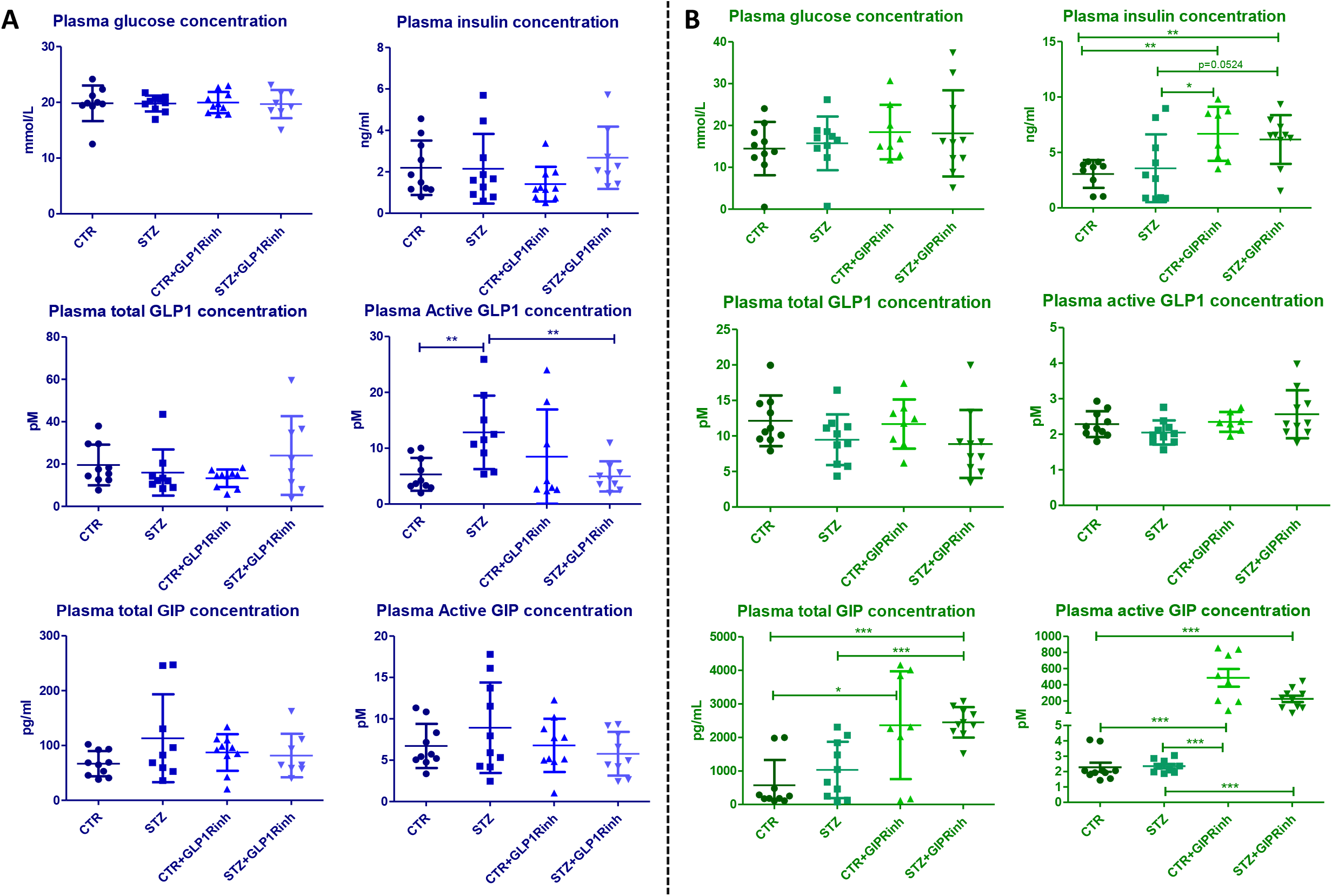
Glucose, insulin, glucagon-like peptide-1, and gastric inhibitory polypeptide plasma concentrations. One month after intracerebroventricular (icv) streptozotocin (STZ) or vehicle (CTR) administration rats were injected icv with 85 µg/kg of glucagon-like peptide-1 receptor antagonist (Exendin fragment 9-39, GLP1Rinh; experiment 1), 85 µg/kg of gastric inhibitory polypeptide receptor antagonist (Pro3-GIP, GIPRinh; experiment 2) dissolved in saline or with saline only (CTR and STZ). Animals were sacrificed 30 minutes after icv administration and blood was sampled for analysis of glucose, insulin, total and active GLP1, total and active GIP concentrations in plasma (**A** and **B**). Values are presented as vertical scatter plots with mean +/- SD and data analyzed by a non-parametric Kruskal-Wallis one-way ANOVA test followed by a Mann Whitney U test (*p<0.05; **p<0.01; ***p<0.001).

### Central GLP-1R inhibition has a higher impact on the brain proteins involved in cell energy

The level of proteins involved in the brain cell energy status was altered region-dependently and more pronounced following central GLP-1R inhibition, particularly in HPT of the STZ group compared to the controls, respectively. Both GLP-1R and GIPR inhibition decreased hypothalamic COXIV signal in the STZ group by 76.2% (p=0.0079) and by 57.7% (p= 0.0087) versus the untreated STZ group, respectively, and GLP-1R inhibition decreased it by 65.5% in STZ compared to the CTR+GLP1Rinh group, respectively (Fig 3A). However, only GIPR inhibition altered the level of COXIV in HPC by increasing it in the control group (+191%, p=0.0519) (Fig 3B). CytC levels remained unchanged after central GIPR and GLP-1R inhibition in both brain regions (Fig 3A and B). PDH levels were altered only after GLP-1R inhibition in both regions; decreased in the HPC of controls (−60.6%, p=0.0022 vs CTR alone) and increased in HPT of the STZ group both versus STZ alone (+66.9%, p=0.0556) and CTR+GLP1Rinh (+96.1%, p=0.0043) group (Fig 3A). PDH levels remained unchanged following GIPR inhibition (Fig 3B).

**Fig. 3.**
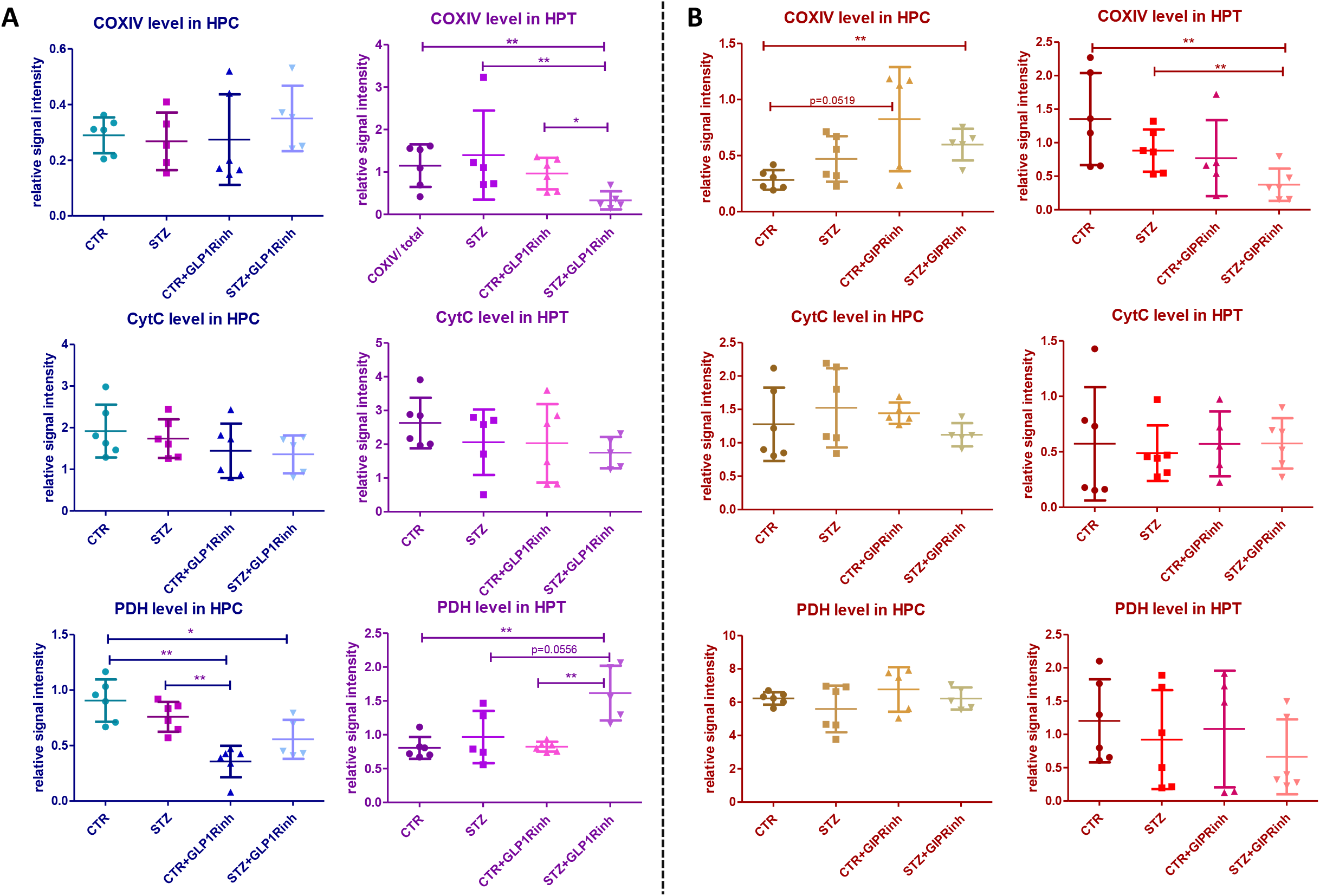
Impact of central glucagon-like peptide-1 and gastric inhibitory polypeptide inhibition on the levels of cytochrome C, cytochrome C oxidase IV, and pyruvate dehydrogenase in the hippocampus and hypothalamus. One month after intracerebroventricular (icv) streptozotocin (STZ) or vehicle (CTR) administration rats were injected icv with 85 µg/kg of glucagon-like peptide-1 receptor antagonist (Exendin fragment 9-39, GLP1Rinh; experiment 1), 85 µg/kg of gastric inhibitory polypeptide receptor antagonist (Pro3-GIP, GIPRinh; experiment 2) dissolved in saline or with saline only (CTR and STZ). Animals were sacrificed 30 minutes after icv administration and the hippocampus (HPC) and hypothalamus (HPT) were dissected, homogenized/sonicated and protein concentration was measured. Cytochrome C (CytC), cytochrome C oxidase IV (COXIV), and pyruvate dehydrogenase (PDH) levels in the HPC and HPT were measured by Western blot analysis 30 minutes after central GLP1R (**A**) and GIPR (**B**) inhibition. Values are presented as vertical scatter plots with mean +/- SD and data analyzed by a non-parametric Kruskal-Wallis one-way ANOVA test followed by a Mann Whitney U test (*p<0.05; **p<0.01; ***p<0.001).

### Central inhibition of GIPR has a more pronounced impact on the AMPK protein level and activity

AMPK plays a significant role in regulating the cell metabolism and energy status; and recent studies indicate that it may play a key role in AD (Assefa et al., 2020). Phosphorylation of AMPK at Thr(172) is important for activation of the kinase (Stein et al., 2000). The change in signal/activity of AMPK was pronounced following GIPR inhibition in a region-dependent manner in the control and STZ group. The level of pAMPK was decreased in HPC of both groups following GIPR inhibition (CTR+GIPRinh vs CTR, - 37.5%, p=0.0087; STZ+GIPRinh vs STZ, -25.9%, p=0.0260), while the increase observed in HPT was insignificant (Fig 4A and B). The level of total AMPK after GIPR inhibition was decreased in HPC of the controls (−59.2%, p=0.0022) and in the STZ alone compared to the CTR alone group (−51.2%, p= 0.0043), while in HPT, a decrement following GIPR inhibition was found in the STZ group (−55% vs STZ alone, p=0.0303; -49.2% vs CTR+GIPRinh, p= 0.0303) (Fig B). AMPK activity, assessed indirectly by the phospho/total AMPK ratio, was found decreased following GIPR inhibition in HPC of the STZ group (−46.3%, p=0.0022 vs STZ; -46%, p=0.0043 vs CTR+GIPRinh) (Fig 4A and B). In contrast to HPC (no change in AMPK activity in STZ alone vs CTR alone groups), in HPT, AMPK activity was found decreased in the STZ alone group compared to the CTR alone one (−24.1%, p=0.0152), while after GIPR inhibition it was significantly increased in the STZ group (+228.2% vs STZ, p=0.0043; +71.1%, p=0.0159 vs CTR+GIPRinh) (Fig 4B). However, the only change observed following GLP-1R inhibition was decreased pAMPK level in HPT of the STZ group (−34% vs STZ alone, p=0.0317; -34%, p=0.0519 vs CTR+GLP1Rinh) and decreased AMPK activity in HPC of the STZ group (−34% vs STZ alone, p=0.0381; -41.6% vs CTR+GLP1Rinh, p=0.0095) (Fig 4A).

**Fig. 4.**
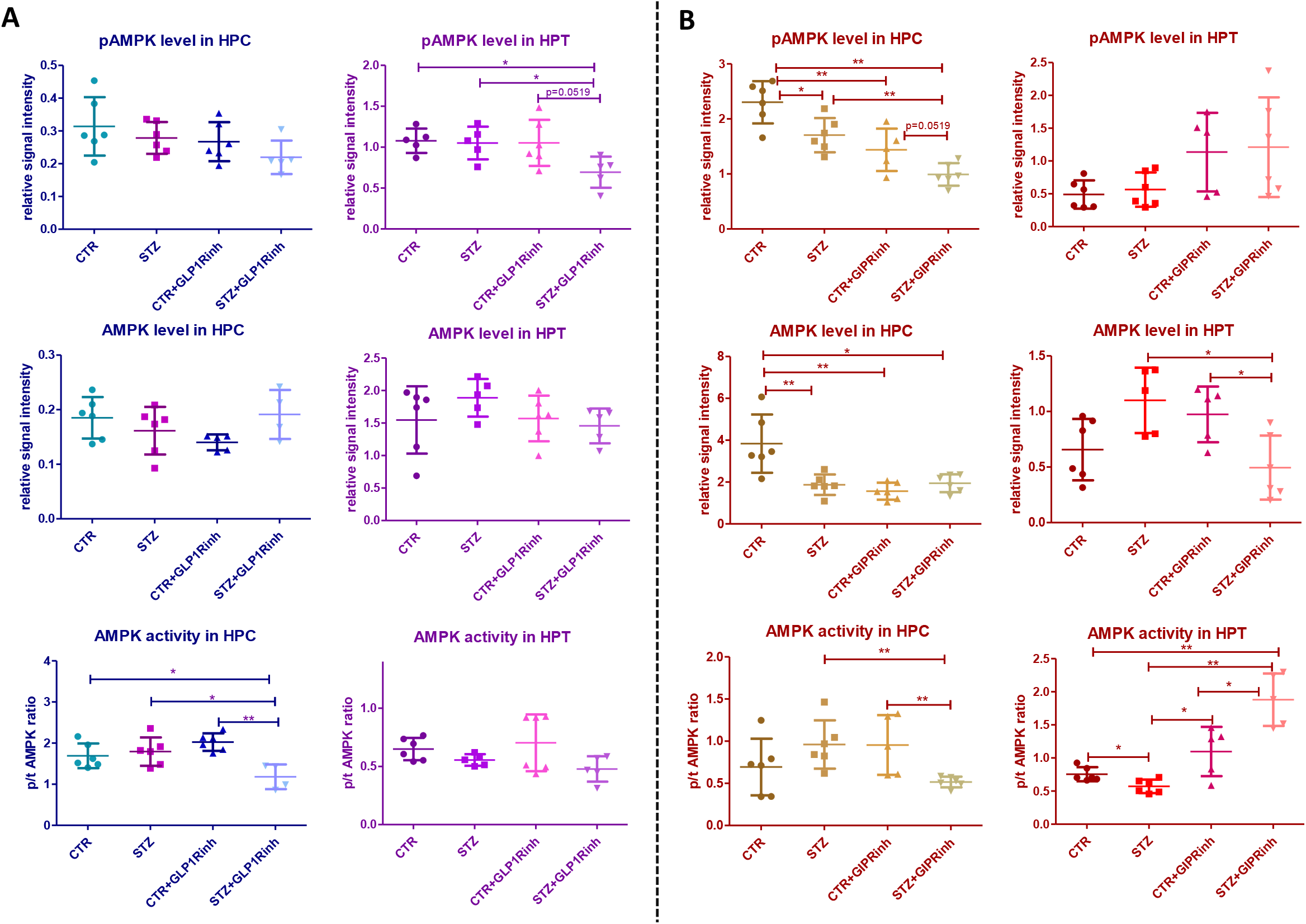
Level and activity of AMP-activated protein kinase in the hippocampus and hypothalamus is changed after central glucagon-like peptide-1 and gastric inhibitory polypeptide inhibition. One month after intracerebroventricular (icv) streptozotocin (STZ) or vehicle (CTR) administration rats were injected icv with 85 µg/kg of glucagon-like peptide-1 receptor antagonist (Exendin fragment 9-39, GLP1Rinh; experiment 1), 85 µg/kg of gastric inhibitory polypeptide receptor antagonist (Pro3-GIP, GIPRinh; experiment 2) dissolved in saline or with saline only (CTR and STZ). Animals were sacrificed 30 minutes after icv administration and the hippocampus (HPC) and hypothalamus (HPT) were dissected, homogenized/sonicated and protein concentration was measured. Phosphorylated (Thr172) and total AMP-activated protein kinase (pAMPK and tAMPK) levels in the HPC and HPT were measured by Western blot analysis 30 minutes after central GLP1R (**A**) and GIPR (**B**) inhibition. Values are presented as vertical scatter plots with mean +/- SD and data analyzed by a non-parametric Kruskal-Wallis one-way ANOVA test followed by a Mann Whitney U test (*p<0.05; **p<0.01; ***p<0.001).

### Central inhibition of GLP-1R affects the level of the neuronal activity marker c-fos, while GIPR inhibition decreases cAMP

Activation of adenylate cyclase and production of cAMP are parts of both the GIP and GLP-1 signaling pathways (Gabe et al., 2020; Marzook et al., 2021) but in our experiments, cAMP was changed only after central GIPR inhibition. Decrement in cAMP level was observed following GIPR inhibition in HPC but not in HPT, being pronounced in healthy rats (−46% vs CTR alone, p=0.0043), and to a much lesser extent in the STZ group (−18.1% vs STZ alone, p=0.0649), although the untreated STZ group had significantly decreased cAMP level compared to the untreated controls (−43.4%, p= 0.0022) (Fig 5B). In contrast, the c-fos level was changed only after GLP-1R inhibition and in both regions; in HPT decreased c-fos was observed in both the healthy (−47.1%, p=0.0411) and STZ group (−49.5%, p=0.0095) while in HPC increased c-fos was measured only in the STZ group (146.6%, p=0.0087 vs STZ alone) (Fig 5A). ATP concentration remained unchanged in both experiments in both brain regions (Fig 5A and B).

**Fig. 5.**
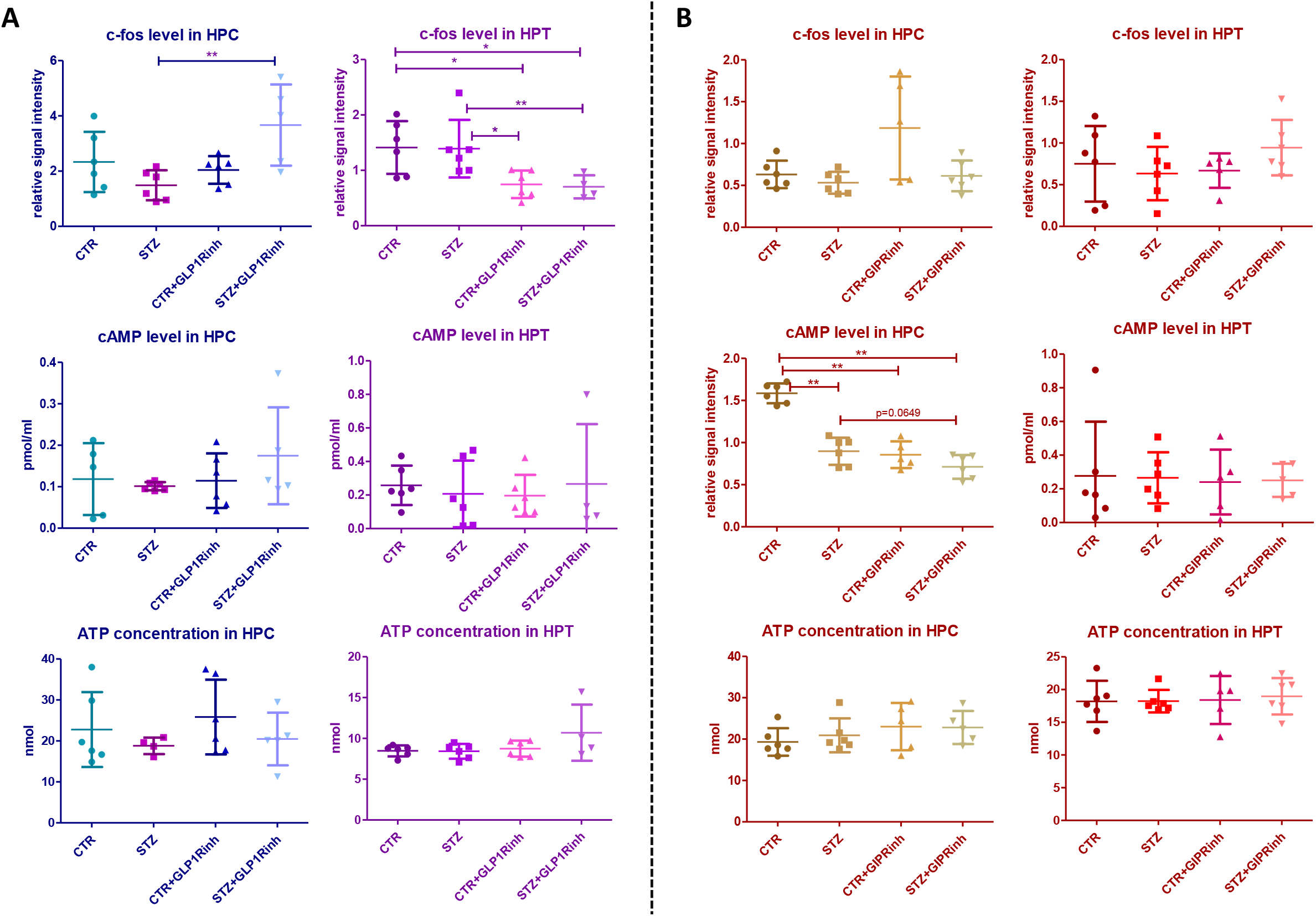
Influence of central glucagon-like peptide-1 and gastric inhibitory polypeptide inhibition on c-fos and cAMP levels and ATP concentration in the hippocampus and hypothalamus. One month after intracerebroventricular (icv) streptozotocin (STZ) or vehicle (CTR) administration rats were injected icv with 85 µg/kg of glucagon-like peptide-1 receptor antagonist (Exendin fragment 9-39, GLP1Rinh), 85 µg/kg of gastric inhibitory polypeptide receptor antagonist (Pro3-GIP, GIPRinh) dissolved in saline or with saline only (CTR and STZ). Animals were sacrificed 30 minutes after icv administration and the hippocampus (HPC) and hypothalamus (HPT) were dissected, homogenized/sonicated and protein concentration was measured. Neuronal activation was measured indirectly through c-fos level by Western blot analysis 30 minutes after central GLP1R (**A**) and GIPR (**B**) inhibition. cAMP levels were measured using a commercial cAMP Direct Immunoassay kit and Western blot analysis. ATP concentration was measured by bioluminescence assay (**A** and **B**). Values are presented as vertical scatter plots with mean +/- SD and data analyzed by a non-parametric Kruskal-Wallis one-way ANOVA test followed by a Mann Whitney U test (*p<0.05; **p<0.01; ***p<0.001).

## Discussion

Our results indicate a substantial impact of central inhibition of incretin receptors with consequences seen in both the periphery and brain, predominantly following the GIPR inhibition, which led to increased plasma active GIP concentration 30 minutes after infusion, while GLP-1R inhibition failed to change plasma GLP-1 levels in healthy rats (Fig 2). Previous reports pointed indirectly to the central regulation of peripheral GIP release based on decreased plasma GIP concentration following insulin icv treatment in a canine model (Yavropoulou et al., 2009) and increased plasma GIP levels following human GIP icv treatment in nonhuman primates (Higgins et al., 2016). Since there is no literature data on the effects of central GIPR inhibition on plasma GIP, increased plasma GIP in our research might seem contradictory to the work of Higgins et al. (Higgins et al., 2016). The reasons for possible inconsistencies could only be speculated; (i) (Pro3)GIP used as GIPR antagonist in our research acts as a weak partial agonist in mice and rats (Sparre-Ulrich et al., 2016); (ii) acute block of the central GIP system induced by a bolus dose of GIPR inhibitor might acutely activate feedback mechanisms leading to increased peripheral GIP secretion, which could be inverted to decreased plasma GIP levels in long-term central GIPR inhibition; (iii) it could not be excluded that to a certain extent, substantially high plasma GIP levels in our research reflects the inability of the 6A1A anti-GIP mouse IgG (used in active GIP ELISA kit) to discriminate between the N-terminal sequence of the GIP(1-42) and [Pro^3^]-GIP; (iv) Higgins et al. observed that GIP given icv can stimulate increased plasma GIP in the absence of nutrient stimulation; however, postprandial GIP levels were not different after icv GIP infusion compared to controls (Higgins et al., 2016) and animals in our research had *ad libitum* access to food; (v) the observed increase in plasma insulin levels might provide indirect support that plasma GIP levels have indeed been increased following acute inhibition of central GIPR but further research should elucidate whether increased plasma insulin is an indirect or direct effect of central GIPR inhibition, or maybe both.

Although direct data on central regulation of plasma GLP-1 levels are lacking, such a possibility seems likely based on the evidence that stimulation of brain GLP-1R in rodents modulates insulin and glucagon secretion, glucose homeostasis, and corticosterone plasma levels (Jessen et al., 2017; Knauf et al., 2005; Sandoval et al., 2008; Tudurí et al., 2015). Modest data on the effects of central GLP-1R inhibition indicates that during hyperglycemia such a treatment leads to increased peripheral glucose utilization and insulin sensitization in mice (Knauf et al., 2005). We have demonstrated that in healthy rats acute inhibition of central GLP-1R (in contrast to GIPR inhibition) did not alter either insulin, glucose, GLP-1 or GIP plasma concentrations (Fig 2). The only change seen 30 min following GLP-1R inhibition in the STZ-icv rat model of sAD is decreased plasma concentration of active GLP-1 (Fig 2A). A lack of this effect in control rats confirms previously observed dysfunctional GLP-1 homeostasis in this sAD model (Knezovic et al., 2018). It could not be excluded that the acute block of GLP-1 signal in the brain following central GLP-1R inhibition is compensated by GLP-1 secretion within the brain occurring in the nucleus of solitary tract (Vrang et al., 2007). Literature data on GIP brain secretion are inconsistent, demonstrating both its presence (Nyberg et al., 2005; Tseng et al., 1993) and absence (Adriaenssens et al., 2020; Kaneko et al., 2019). The observed discrepancy between experiments could be due to experiments not being conducted simultaneously and using different rat litters. Unfortunately, this can only be circumvented by conducting the experiments at the same time with the same rat litter, which is sometimes impossible because of the large number of animals needed.

Although both GIP and GLP-1 act like insulin secretagogues in physiological conditions, insulin secretion in T2DM can be stimulated by supraphysiological doses of GLP-1 but not GIP (Nauck et al., 1993). Neurodegenerative disorders, AD in particular, are associated with T2DM-like metabolic abnormalities and insulin resistance in the brain, due to which intranasal insulin, insulin sensitizers and GLP-1R agonists have been considered as potential therapeutic options in AD treatment (Kellar and Craft, 2020; Watson and Craft, 2003). Like diabetes, obesity also contributes to the development of AD and studies have identified several overlapping mechanisms of these disorders (Pugazhenthi et al., 2017). Basic research indicates more efficient neuroprotection of neurodegenerative and AD hallmarks with dual GLP-1/GIP agonists than GLP-1 analogues alone (Ji et al., 2016). Additionally, GIPR and GLP-1R knockout mice showed impairment of synaptic plasticity and cognitive deficit (Abbas et al., 2009; Faivre et al., 2011; Lennox et al., 2015); however, in another research, GIPR knockout mice showed extended lifespan, increased exploratory and decreased anxiety-based behaviors (Hoizumi et al., 2019), indicating a divergent effect of incretins.

Efflux of inhibitors into the periphery following intracerebroventricular administration should also be acknowledged as an important limitation of the presented research. We have mentioned in our recent research (Homolak et al., 2021) a solution presented by Kanoski et al. They introduced another group that received an intraperitoneal GLP-1R inhibitor in doses that could have effluxed after icv injection and concluded that this ip treatment did not significantly attenuate intake suppression by ip delivery of agonists (the icv inhibitor antagonized peripherally administered liraglutide and exendin-4 that accessed the brain and acted directly on the CNS GLP-1R) (Kanoski et al., 2011). In future research, introducing another group with peripherally administered inhibitors can be added to further distinguish central and peripheral effects.

Our results demonstrate that GIP is more important for brain cell energy and metabolism than GLP-1 and thus support and provide a possible explanation for better neuroprotection of dual agonists. Direct inhibition of brain GIPR is associated with a different pattern of changes in HPC compared to HPT, seen as the opposite direction of changes of the proteins involved in metabolism and energy (COXIV and AMPK, Fig 3B and 4B). AMPK acts as a sensor of cellular energy and it is activated in response to falling energy charge by stimulating energy production via catabolic pathways, while decreasing nonessential energy-consuming pathways to restore cellular energy stores (O’Neill, 2013; O’Neill et al., 2013). Changes in AMPK homeostasis were predominantly seen following GIPR inhibition, demonstrating a region- and group-dependent effect with AMPK activity found decreased in the STZ group in HPC and increased in HPT, while in controls decreased protein levels of phospho and total AMPK (but not the activity) were found in HPC only. Central GLP-1R inhibition decreased AMPK activity in HPC only in the STZ group. To this date, there is no data on the effect of central GIPR modulation on AMPK levels and activity in the brain. Since GIP and GLP-1 are involved in glucose homeostasis, further research on the relationship between hypothalamic AMPK (considered as a regulator of the whole-body energy balance) and GIP/GLP-1 signaling could give insight into possible new therapeutic approaches for the treatment of obesity, diabetes or neurodegenerative disorders. Unchanged ATP concentration in HPC and HPT (Fig 5) could be due to the activation of compensatory mechanisms since it was measured 30 minutes after inhibition and the ATP turnover rate is high.

Recently, novel therapeutics targeting GIP signaling, GIP antagonists, and dual GIPR and GLP-1R agonists regain interest in the GIP regulation of energy homeostasis in the brain (Adriaenssens et al., 2020). Mitochondrial COX is the primary site of cellular oxygen consumption and is essential for aerobic energy generation in the form of ATP (Timón-Gómez et al., 2018). Like cytochrome C, it is used as a mitochondrial marker. The opposite pattern of protein levels involved in energy in HPC and HPT after GIPR inhibition indicates a divergent action of GIP in these regions. Possible cell energy deprivation in HPT (presented as decrement of COXIV; Fig 3; and increment of AMPK activity; Fig 4) can lead to increased peripheral secretion of GIP and insulin, and opposite changes of protein levels (presented as COXIV level increment; Fig 3; and AMPK level decrement; Fig 4) found in HPC can give insight into the possible involvement of GIPR signaling in cognition. Further research is needed to elucidate the action of central GIPR inhibition on learning and memory.

The majority of changes seen after acute central GLP-1R inhibition were seen in animals that were administered STZ-icv 1 month before, suggesting that STZ induces an imbalance in the central GLP-1 system. As previously reported, the therapeutic window for the efficiency of different drug treatments is up to 3 months after STZ-icv administration, a time-point when cognitive decline in this model enters into the dose-dependent progressive and irreversible phase, while 1 month after STZ-icv treatment some reversibility of the pathology could still be observed (Knezovic et al., 2015; Salkovic-Petrisic et al., 2013). How GLP1 system is involved in STZ induced pathophysiological changes or development of AD it is still not yet clear and it needs further elucidation. Most of the research regarding GLP1 system in AD is based on testing the agonists or DPPIV inhibitors as potential neuroprotective agents and strategy to normalize insulin sensitivity in the brain (Markaki et al., 2020). The potential involvement of GLP1 signaling in pathogenesis of human sAD has not yet been fully explored and it needs further evaluation of possible involvement in neurodegeneration. In pancreatic β-cells, chronic stimulation of GLP-1R increases glycolysis and ATP production through transcriptional activation and expression of glycolytic genes, and inhibition of PI3K/mTOR pathway abolishes such an effect (Carlessi et al., 2017). These findings are in concordance with a decreased level of PDH in HPC after GLP-1R inhibition, suggesting decreased glucose metabolism. If GLP-1R inhibition decreases the import of glucose, PDH activity is downregulated to limit the use of glucose by oxidative phosphorylation (Milne, 2013). In contrast to the results found in HPC, PDH levels in HPT were increased in STZ rats, in compliance with decreased COXIV level after GLP-1R inhibition. Whether an increased PDH level is a compensatory effect due to decreased COXIV or vice versa, remains to be explored. In contrast, GIPR inhibition did not alter PDH levels at all, further suggesting different actions of GIP and GLP-1 in the brain (Fig 3). In line with these results, only GLP-1R inhibition influenced neuronal activation, increasing it in HPC and decreasing it in HPT. Changes found in proteins involved in cell energy seemed not to have any influence on the marker of neuronal activity.

Due to highly increased GIP levels in plasma after acute central GIPR inhibition and the unchanged plasma GLP-1 levels after central GLP-1R inhibition, we can presume that the CNS is more dependent on the production of GIP at the periphery than GLP-1. Based on literature data (Fu et al., 2020), it seems more likely that a signal to the periphery for the elevation of GIP secretion might originate from the hypothalamus than from the hippocampus, but further research is needed to confirm this hypothesis. Changes observed after GLP-1R inhibition mostly concern STZ-icv rats, and to a much lesser extent healthy controls, further supporting our previously published data on dysfunctional GLP-1 signaling in the STZ-icv rat model of sAD and involvement of the brain-gut GLP-1 axis in the maintenance of redox balance in the upper small intestine (Homolak et al., 2021). Data presented here indicates that GIPR and GLP-1R inhibition, respectively, have the opposite effect on the level of proteins involved in metabolism and energy balance regulation in the hippocampus and hypothalamus. Brain incretin signaling has emerged as a new field that might provide valuable contributions to the research of both the pathophysiology and, accordingly, novel therapeutic strategies in the treatment of diabetes, obesity, aging, and related neurodegenerative disorders.

## Acknowledgement

This work was funded by the Croatian Science Foundation (IP-2018-01-8938 and IP-2014-09-4639). Research was co-financed by the Scientific Centre of Excellence for Basic, Clinical and Translational Neuroscience (project “Experimental and clinical research of hypoxic-ischemic damage in perinatal and adult brain”; GA KK01.1.1.01.0007 funded by the European Union through the European Regional Development Fund).

## Author Contributions

AK and JOB equally contributed to the paper. AK, JOB, JH and ABP performed in vivo part of this research. AK and JOB conducted all assays, ELISA and Western blot measurements in plasma, HPC and HPT. AK performed data analysis and visualization. Both AK and JOB wrote the manuscript. MSP, PI of the lab, contributed to the design of the experiments and supervised the project. JH, ABP and MSP revised the manuscript and provided critical input. All authors have agreed to the published version of the manuscript.

## Materials and Methods

### Animals

Adult (3-month-old) male Wistar rats weighing 250-350 g (Department of Pharmacology, University of Zagreb School of Medicine) were used in the experiment. All animals were housed in cages (2-3 rats per cage) in the animal facility at the Department, kept on standardized food pellets and water *ad libitum*, and maintained under a 12/12 h light/dark cycle.

### Ethics

The experiments were carried out in compliance with current institutional (University of Zagreb School of Medicine), national (Animal Protection Act, NN 102/17), and international (Directive 2010/63/EU) guidelines on the use of experimental animals. The national regulatory body, the Croatian Ministry of Agriculture, approved the experiments (license number EP 186 /2018).

### Materials

Streptozotocin, Exendin fragment 9-39, PhosSTOP phosphatase inhibitor tablets, and protease inhibitor cocktail were purchased from Sigma-Aldrich (St. Louis, Missouri, USA). [Pro3]-GIP (Rat) was purchased from Tocris, a Bio-Techne brand (Abingdon, United Kingdom). The glucose measuring kit (Greiner Diagnostic Glucose GOD-PAP) was acquired from Dijagnostika (Sisak, Croatia). The chemiluminescent Western blot detection kit (SuperSignal West Femto Chemiluminescent Substrate) and ATP determination kit were from Thermo Scientific (Rockford, IL, USA). The ELISA Kit for rat/mouse insulin, GLP1 and GIP total ELISA kits, and high-sensitivity GLP1 Active ELISA kit were acquired from Merck Millipore (Billerica, USA). Polyclonal anti-c-Fos antibody produced in rabbit was purchased from Abcam (Cambridge, UK). TGX FastCast Acrylamide Solution was purchased from Bio-Rad (Hercules, California, USA). Monoclonal anti-phospho-AMPKα (Thr172) produced in rabbit, polyclonal anti-AMPKα produced in rabbit, monoclonal mouse anti-cytochrome c oxidase subunit 4 (COXIV), monoclonal rabbit anti-cytochrome c (CytC), monoclonal rabbit anti pyruvate dehydrogenase (PDH), anti-mouse IgG horseradish peroxidase-linked antibody, anti-rabbit IgG horseradish peroxidase-linked antibody were acquired from CellSignaling (Beverly, MA, USA). cAMP Direct Immunoassay Kit was purchased from BioVision (Milpitas, California, USA). Polyclonal rabbit anti-cAMP antibody (Elabscience, Houston, Texas, USA) was purchased from antibodies-online. Mouse GIP active form, high sensitivity, ELISA kit was purchased from Tecan IBL International (Hamburg, Germany).

### Experimental design

#### Streptozotocin treatment

Two experiments with 4 groups each were conducted with the same procedure. Three-month-old male Wistar rats were subjected to general anesthesia (ketamine 70 mg/kg; 7 mg/kg xylazine), followed by intracerebroventricular injection of STZ (3 mg/kg, dissolved in 0.05 M citrate buffer, pH 4.5, bilaterally 2 µL/ventricle, split into two doses administered on day 1 and 3), according to the procedure first described by Noble et al. (Noble et al., 1967) and used in our previous experiments (Babic Perhoc et al., 2019; Grünblatt et al., 2007; Knezovic et al., 2015; Osmanovic Barilar et al., 2015; Salkovic-Petrisic et al., 2014). Control (CTR) animals were given an equal volume of vehicle (0.05M citrate buffer, pH 4.5) icv by the same procedure on days 1 and 3 (Fig 1).

#### Experiment 1: GLP1Rinh treatment

Control and STZ-treated rats were randomly divided in four groups. Two groups (CTR+GLP1Rinh and STZ+GLP1Rinh) received icv 85 µg/kg (25.23 mmol/kg) of the competitive GLP-1 receptor antagonist (Exendin fragment 9-39) dissolved in saline (bilaterally 2 µL/ventricle), administration protocol based on our preliminary experiment. The other 2 groups (CTR and STZ) received saline only (bilaterally 2 µL/ventricle). All animals were sacrificed 30 minutes after icv administration. There were 10 animals per group, except 9 animals in the STZ+GLP1Rinh group (Fig 1).

#### Experiment 2: GIPRinh treatment

Control and STZ-treated rats were randomly divided in four groups. Two groups (CTR+GIPRinh and STZ+GIPRinh) received icv 85 µg/kg (12.19 mmol/kg) of GIP receptor antagonist (Pro3-GIP) dissolved in saline (bilaterally 2 µL/ventricle). The other 2 groups (CTR and STZ) received saline only (bilaterally 2 µL/ventricle). All animals were sacrificed 30 minutes after icv administration. There were 10 animals per group, except 8 animals in the CTR+GIPRinh group (Fig 1).

### Tissue preparation and blood sampling

Sacrification was performed under deep general anesthesia with thiopental and diazepam (60 mg/kg and 6 mg/kg). Blood was sampled from retro-orbital sinus in tubes containing heparin and DPP-4 inhibitor and centrifuged for 10 min at 3000 rpm at 4°C; supernatant (plasma) was collected and stored at -80°C. Six or five animals per group were decapitated after which brains were quickly removed, the hippocampus (HPC) and hypothalamus (HPT) dissected out and frozen in liquid nitrogen, and stored at -80°C. Brain tissue samples for protein analysis were thawed and homogenized/sonicated with four volumes of lysis buffer containing 10mM HEPES, 1mM EDTA, 100 mM KCl, 1% Triton X-100, pH 7,5, protease inhibitor cocktail (1:100), and phosphatase inhibitor tablets. Homogenates were centrifuged at 13000 rpm for 10 min at 4°C and the supernatant was frozen and stored at -80°C. Protein concentration was measured by Lowry protein assay.

### Biochemical analysis

#### Glucose measurements

Quantitative determination of glucose concentration in plasma was done by enzymatic colorimetric method (GOD/PAP) using a commercial kit. The measurement was conducted by mixing 2 µL of rat plasma samples and 200 µL of reagent and absorbance was measured using the microplate reader Infinite 200 PRO (Tecan Trading AG, Switzerland) after 10 minutes at 500 nm. The concentration of glucose was expressed in mmol/L (mM).

#### Insulin measurements

Plasma insulin levels were measured using a commercial rat/mouse insulin ELISA kit. Rat plasma samples (10 µL) were added to appropriate wells. The assay procedure was conducted by adhering to the manufacturer’s protocol. Absorbance was measured at 450 nm, subtracted for the absorbance at 590 nm using a microplate reader (Infinite 200 PRO) and the difference of absorbance was recorded. The concentration of insulin was expressed in ng/mL.

#### GLP-1 total measurements

Plasma GLP-1 total levels were measured using a commercial multi-species GLP-1 total ELISA kit. Rat plasma samples were added in a volume of 50 µL to appropriate wells. The assay procedure was conducted by adhering to the manufacturer’s protocol. Absorbance was measured at 450 nm and 590 nm using a microplate reader (Infinite 200 PRO) and the difference of absorbance was recorded. The concentration of GLP-1 total was expressed in pM.

#### GIP total measurements

Plasma GIP total levels were measured using a commercial rat/mouse GIP (total) ELISA kit. Rat plasma samples were added in a volume of 10 µL to appropriate wells. The assay procedure was conducted by adhering to the manufacturer’s protocol. Absorbance was measured at 450 nm and 590 nm using a microplate reader (Infinite 200 PRO) and the difference of absorbance was recorded. The concentration of GIP was expressed in pg/mL.

#### GLP-1 active measurements

Plasma GLP-1 active levels were measured using a commercial high-sensitivity GLP-1 active ELISA kit. Test samples (50 µL), undiluted, were added to the appropriate wells. Measurement of relative light units was done in a luminometer, the GloMax microplate reader (Promega, Madison, USA) within 5 minutes after adding the substrate solution. The concentration of GLP-1, active form, was expressed in pM.

#### GIP active measurements

Plasma GIP active levels were measured using a commercial GIP active ELISA kit. Briefly, samples were thawed and diluted with a buffer from the kit. Based on preliminary measurements, plasma from the animals that have received GIPRinh was diluted 5 times, the other groups were not diluted at all. In total 100 µL of test samples were added to the wells. The measurement was conducted by adhering to the manufacturer’s procedure. Absorbance was measured at 450 nm and sub-wavelength at 620 nm using the microplate reader Infinite 200 PRO. The concentration of GIP active was expressed in pM.

#### ATP measurements

Quantitative determination of ATP concentration in tissue homogenates (HPC and HPT) was done by bioluminescence assay with recombinant firefly luciferase and its substrate D-luciferin using a commercial kit. The measurement was conducted by mixing 10 µL of rat tissue homogenates and 100 µL of standard reaction solution reagent and luminescence was measured in luminometer, the GloMax microplate reader, and background luminescence was subtracted and the amount of ATP was calculated from the standard curve. The concentration of ATP was expressed in nM.

#### cAMP measurements

Rat brain tissue (HPC and HPT) cAMP levels were measured using a commercial cAMP Direct Immunoassay kit. Briefly, samples and standards were acetylated at room temperature for 10 min, then 50 µL of the acetylated samples and standards were added to each well. The measurement was conducted by adhering to the manufacturer’s procedure. Absorbance was measured at 450 nm using the Infinte 200 PRO microplate reader. The concentration of cAMP was expressed in pmol.

#### Western blot analysis

Equal amounts of total protein (35 µg per sample) in the HPT and HPC were separated by sodium dodecyl sulfate-polyacrylamide gel electrophoresis using TGX Stain-Free 12% gels (gels were visualized using ChemiDoc Imaging Systems, Bio-rad, Hercules, California, USA) and transferred to nitrocellulose membranes using Trans-Blot Turbo Transfer System (Bio-rad, Hercules, California, USA). Transfer efficiency was checked using the same stain-free system and 0.1% Ponceau S in 5% acetic acid solution. The nitrocellulose membranes were blocked for 1 h at RT in 5% non-fat milk, added to low-salt washing buffer (LSWB) containing 10 mM Tris, 150 mM NaCl, pH 7.5, and 0.5% Tween 20. The blocked membranes were incubated with primary anti-CytC (1:1000), anti-COXIV (1:1000), anti-PDH (1:1000), anti-cAMP (1:2000), anti-c-fos (1:2000), anti-phospho-AMPK (1:1000) and anti-total-AMPK antibodies (1:1000) overnight at 4°C. After incubation, the membranes were washed 3x with LSWB and incubated for 1 h at RT with secondary antibody solution (anti-rabbit or anti-mouse IgG, 1:2000). After 3x washing in LSWB, signals were visualized using a chemiluminescence Western blotting detection reagent. The signals were captured and visualized with a MicroChemi video camera system (DNR Bio-Imaging Systems). Proteins were analyzed using ImageJ software and blots were normalized to total protein signal of UV-transilluminated gels using the same analysis procedure.

### Statistics

Data were presented as vertical scatter plot with mean ± SD with the significance of between-group differences in all analyses tested by two-tailed Kruskal-Wallis one-way ANOVA analysis of variance, followed by Mann-Whitney U-test, with p<0.05 considered statistically significant using GraphPad Prism 5 statistical software.

